# Development of Optically-Controlled “Living Electrodes” with Long-Projecting Axon Tracts for a Synaptic Brain-Machine Interface

**DOI:** 10.1101/333526

**Authors:** Dayo O. Adewole, Laura A. Struzyna, James P. Harris, Ashley D. Nemes, Justin C. Burrell, Dmitriy Petrov, Reuben H. Kraft, H. Isaac Chen, Mijail D. Serruya, John A. Wolf, D. Kacy Cullen

## Abstract

Achievements in intracortical neural interfaces are compromised by limitations in specificity and long-term performance. A biological intermediary between devices and the brain may offer improved specificity and longevity through natural synaptic integration with deep neural circuitry, while being accessible on the brain surface for optical read-out/control. Accordingly, we have developed the first “living electrodes” comprised of implantable axonal tracts protected within soft hydrogel cylinders for the biologically-mediated monitoring/modulation of brain activity. Here we demonstrate the controlled fabrication, rapid axonal outgrowth, reproducible cytoarchitecture, and simultaneous optical stimulation and recording of neuronal activity within these engineered constructs *in vitro*. We also present their transplantation, survival, integration, and optical recording in rat cortex *in vivo* as a proof-of-concept for this neural interface paradigm. The creation and functional validation of these preformed, axon-based “living electrodes” is a critical step towards developing a new class of biohybrid neural interfaces to probe and modulate native circuitry.

## Introduction

Techniques for neuromodulation (such as deep brain stimulation for Parkinson’s disease) and neural recording (commonly called brain-computer interfaces (BCIs)) can electrically stimulate or capture neuronal activity within the brain ^1^. These methods have been developed for a range of investigative and clinical goals, from cochlear implants for the hearing-impaired to computer control for those with neuromuscular disorders ^1^. Despite significant milestones to date, several issues have limited the potential medical impact of these neural interface technologies. Broadly, implantable BCIs use inorganic microelectrodes, which often exhibit diminished performance over time due to biotic (inflammation, neuronal loss, and glial scarring) and abiotic (biostability issues including decreasing impedance due to loss of insulation, mechanical failure) factors, impeding recording quality^1–5^. In neuromodulation, effectiveness is limited by the inability to target specific neurons or neuronal subtypes (e.g. excitatory vs. inhibitory neurons) within the volume of charge injection and the thresholds for both safe and functional therapeutic stimulation ^6^. Specific cell types may be targeted by using optogenetics to activate genetically modified neurons via spatial distribution of light. However, the light scattering properties of tissue block precise photostimulation of neurons more than a few hundred microns deep^7^. Implantable optical fibers, lenses, or micro-LEDs have been used, yet chronic performance is limited by the foreign body response and/or overheating of surrounding tissue^8,9^. Further, the longevity and immune response in humans is unknown for virally transduced optogenetic proteins. Finally, across electric and/or optical input-output paradigms, the information transfer bandwidth limits the quality of the neural interface. The ability to address these design challenges – compatibility with the brain, target specificity, and long-term stability – will direct the utility and clinical translation of future neuromodulation and neural recording technologies.

To begin to address these limitations, we have developed micro-Tissue Engineered Neural Networks (μTENNs). μTENNs are comprised of discrete population(s) of neurons connected by long bundles of axons protected within a microscopic hydrogel cylinder (“microcolumn”) (Figure 1)^10,11^. μTENNs were originally developed to reconstruct lost or damaged neuroanatomy following brain injury, and previously demonstrated neuronal survival, maintenance of axonal architecture, and synaptic integration with host cortical neurons following targeted microinjection into rats^10,11^. Here, we further develop this tissue engineering approach into a putative “living electrode”; that is, a self-contained, implantable, synaptically-based conduit to affect neuronal activity. ^10–12^. Our efforts to advance the μTENN technology in this manner may uniquely address challenges in current neuromodulation and neural recording strategies (Figure 1J). In this radical paradigm, the μTENN is implanted at a predetermined depth to synaptically integrate with local neural circuitry and propagate neuronal activity along μTENN axons to/from an externalized apparatus at the brain surface (Figure 1J). The segregation of biological and inorganic material may ameliorate the foreign body response and improve long-term stability. Moreover, the μTENN neuronal phenotype may be selected to preferentially synapse certain neurons or elicit a desired excitatory, inhibitory, or modulatory effect. Finally, μTENNs may be transduced to express optogenetic proteins prior to implant, enabling light-driven neuromodulation (through photostimulation of the μTENN neurons to influence downstream cortical activity) or monitoring (recording μTENN neurons as a representation of multiple cortical synaptic inputs). In this way μTENNs may serve as a biologically-based, selective, and potentially permanent neural interface. The present work details the fabrication and characterization of next-generation axon-based μTENNs *in vitro*, including growth, viability, maturation, and structure. We also demonstrate light-driven control of μTENNs *in vitro*, μTENN implantation, survival, and integration in the rodent cortex, and optical monitoring of μTENN activity *in vivo* as proof-of-concept for optically-controlled living electrodes.

**Figure 1:**
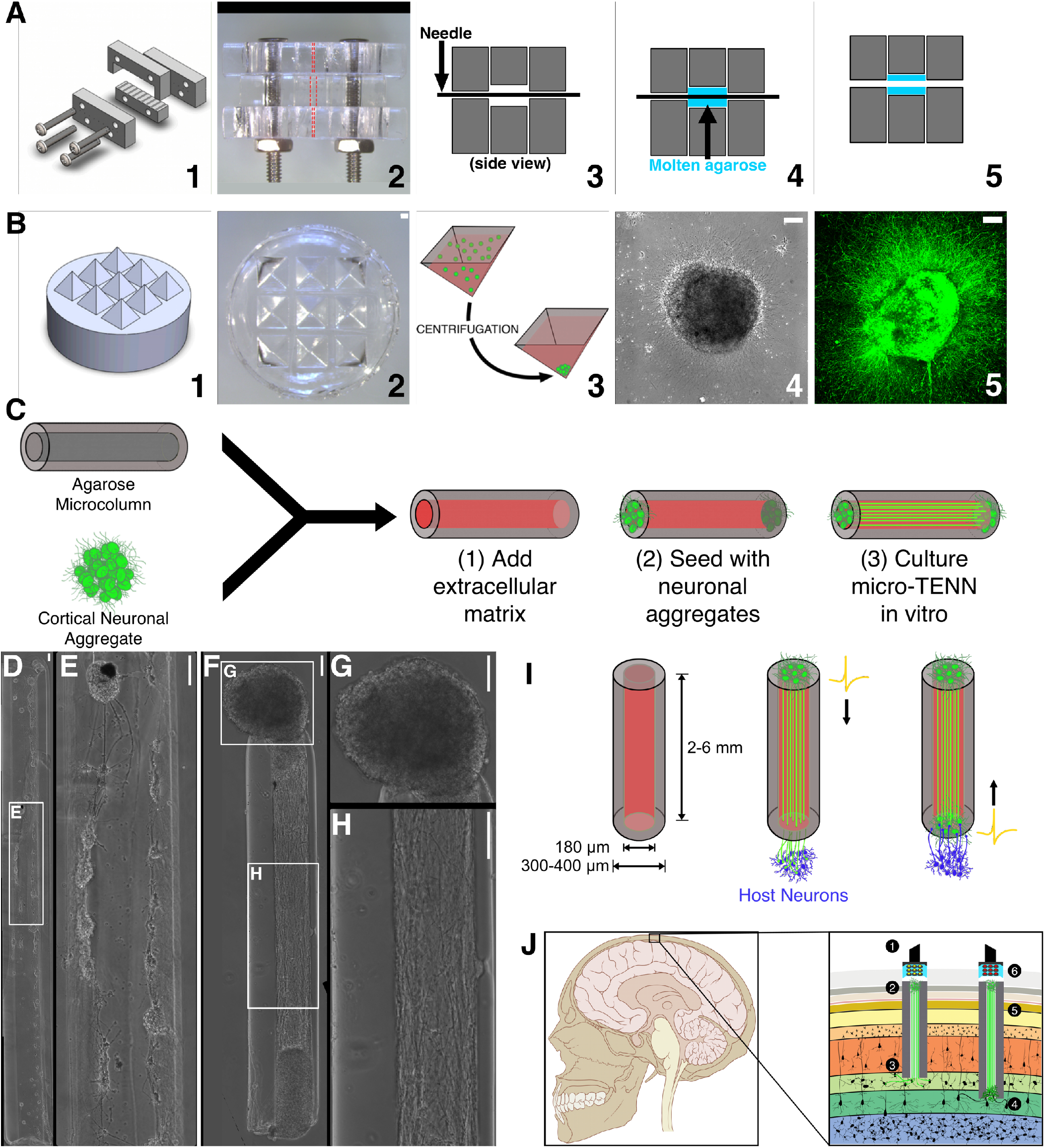
μTENN Fabrication and Living Electrode Concept. μTENNs comprise a hydrogel microcolumn, living neuronal aggregates, and an extracellular matrix lumen. **(A) 1**: A customizable acrylic mold for generating microcolumns. **2**: Top view of the mold (dashed lines represent outer and inner diameters). **3**: Needles of the desired inner diameter are inserted into the mold. **4**: Molten agarose (blue) is introduced into the mold and cooled. **5**: Microcolumns are removed after needle removal/mold disassembly. **(B) 1**: A 3D-printed mold for square pyramidal wells. **2**: Pyramidal wells cast in PDMS. **3**: Dissociated neurons (green) forced into aggregates through centrifugation. **4**: Phase image of an aggregate 24 hours after plating. **5**: Confocal reconstruction of aggregate at 72 hours, labeled with GFP. **(C)** Microcolumns (gray) are filled with an extracellular collagen-laminin matrix (red). Neuronal aggregates are then placed at one or both ends of the microcolumn and grown *in vitro*. **(D)** μTENNs were originally fabricated with dissociated neurons, yielding limited control over axonal growth and network formation **(E)**. Aggregate μTENNs **(F)** exhibit robust axonal growth and more controllable architecture, with discrete regions of cell bodies **(G)** and neuritic projections **(H)**. **(I)** <u>Left</u>: Current μTENN dimensions. <u>Middle</u>: Unidirectional μTENNs synapse host neurons (purple) to relay inputs to targeted cortical regions. <u>Right</u>: Host neurons synapse bidirectional μTENNs, relaying activity from host cortex to the dorsal aggregate. **(J)** μTENNs as transplantable input/output channels. *Inputs:* an LED array (1) optically stimulates a unidirectional, channelrhodopsin-positive μTENN (2) to activate Layer IV neurons (3). *Outputs:* Layer V neurons (4) synapse a bidirectional μTENN (5); relayed neuronal activity is recorded by a microelectrode array (6). Scale bars: 100 μm.

## Materials and Methods

All procedures were approved by the Institutional Animal Care and Use Committees at the University of Pennsylvania and the Michael J. Crescenz Veterans Affairs Medical Center and adhered to the guidelines set forth in the NIH Public Health Service Policy on Humane Care and Use of Laboratory Animals (2015).

### Cortical Neuron Isolation and Culture

Neural cell isolation and culture protocols are similar to that of published work^10,11^. Briefly, timed-pregnant rats were euthanized, and the uterus removed. Embryonic day 18 fetuses were transferred from the uterus to cold HBSS, wherein the brains were extracted and the cerebral cortical hemispheres isolated under a stereoscope via microdissection. Cortical tissue was dissociated in 0.25% trypsin + 1mM EDTA at 37°C, after which the trypsin/EDTA was removed and replaced with 0.15 mg/ml DNase in HBSS. Dissociated tissue + DNase was centrifuged for 3 min at 3000 RPM before the DNase was removed and the cells re-suspended in neuronal culture media, composed of Neurobasal^®^ + B27^®^ + Glutamax™ (ThermoFisher) and 1% penicillin-streptomycin.

### Micro-Tissue Engineered Neural Network (μTENN) Fabrication

μTENNs were constructed in a three-phase process (Figure 1A-C). First, agarose microcolumns of a specified geometry (outer diameter (OD), inner diameter (ID), and length) were formed in a custom-designed acrylic mold as described in earlier work (Figure 1A)^13^. The mold is an array of cylindrical channels that allow for the insertion of acupuncture needles (Seirin, Weymouth, MA) such that the needles are concentrically aligned within the channels. The mold has been fabricated with more precise machining equipment relative to earlier work to better ensure concentric tolerance of the needles and channels. Molten agarose in Dulbecco’s phosphate buffered saline (DPBS) was poured into the mold-needle assembly and allowed to cool (agarose: 3% weight/volume). Once the agarose solidified, the needles were removed and the mold disassembled, yielding hollow agarose microcolumns with a specific outer diameter equal to the size of the channels and inner diameter equal to the outer diameter of the needles. Microcolumns were sterilized via UV light for 30 min and stored in DPBS to prevent dehydration until needed. For these studies, the mold channels were 398 μm in diameter and the acupuncture needles were 180 μm, resulting in microcolumns with a 398 μm OD and a 180 μm ID. Microcolumns were cut to the desired length for each cohort, as described below.

Next, primary cortical neurons were forced into spheroidal aggregates (Figure 1C). These aggregates provide the necessary architecture for the growth of long axonal fascicles spanning the length of the microcolumn. To create aggregates, dissociated cells were transferred to an array of inverted pyramidal wells made in PDMS (Sylguard 184, Dow Corning) cast from a custom-designed, 3D printed mold (Figure 1B). Dissociated cortical neurons were suspended at a density of 1.0-2.0 million cells/ml and centrifuged in the wells at 200g for 5 min. This centrifugation resulted in forced aggregation of neurons with precise control of the number of neurons per aggregate/sphere (12 μL cell suspension per well). Pyramidal wells and forced aggregation protocols were adapted from Ungrin et al ^14^.

Finally, microcolumns were removed from DPBS and excess DPBS removed from the microcolumn channels via micropipette. Microcolumns were then filled with extracellular matrix (ECM) comprised of 1.0 mg/ml rat tail collagen + 1.0 mg/ml mouse laminin (Reagent Proteins, San Diego, CA) (Figure 1C). Unidirectional or bidirectional μTENNs were seeded by carefully placing an aggregate at one or both ends of the microcolumns, respectively, using fine forceps under a stereoscope, and were allowed to adhere for 45 min at 37°C, 5% CO_2_. To create dissociated μTENNs, dissociated cortical neurons were transferred via micropipette into the ECM-filled microcolumn as detailed in prior work ^10,11^. μTENNs were then allowed to grow in neuronal culture media with fresh media replacements every 2 days *in vitro* (DIV).

### Growth Characterization

Phase-contrast microscopy images of μTENNs in culture were taken at 1, 3, 5, 8, and 10 DIV at 10x magnification using a Nikon Eclipse Ti-S microscope, paired with a QIClick camera and NIS Elements BR 4.13.00. μTENNs were fabricated for classification into one of the following groups: dissociated/2 mm long (LE_DISS,2_) (n = 7), unidirectional/2 mm long (LE_UNI,2_) (n = 6), unidirectional/5 mm long (LE_UNI,5_) (n = 3), bidirectional/2 mm long (LE_BI,2_) (n = 15), bidirectional/3 mm long (LE_BI,3_) (n = 12), bidirectional/5 mm long (LE_BI,5_) (n = 17), bidirectional/7 mm long (LE_BI,7_) (n = 8), bidirectional/9 mm long (LE_BI,9_) (n = 3). Growth rates for each group at specific timepoints were quantified as the change in the length of the longest identifiable neurite divided by the number of days between the current and preceding timepoint. The longest neurites were manually identified within each phase image using ImageJ (National Institutes of Health, MD), and length was measured from the edge of the source aggregate to the neurite tip. To standardize measurements, the edge of the source aggregate identified at 1 DIV was used as the reference point across subsequent timepoints. Growth was measured until axons crossed the length of the column (for unidirectional constructs) or axons crossed the distance between aggregates (for bidirectional constructs). Growth rates were averaged for each timepoint, with the average maximum and minimum growth rates and average crossing time compared across aggregate μTENNs with one-way analysis of variance (ANOVA). Post-hoc analysis was performed where necessary with the Bonferroni procedure (p<0.05 required for significance). For reference, planar cultures of cortical neurons (n = 10) were grown in parallel with μTENN cultures, with the longest identifiable neurites measured at 1, 3, and 5 DIV. Single neurites could not be identified at later timepoints due to culture maturation. Axonal outgrowth in planar cultures was taken as the average growth rate across timepoints, which were compared via unpaired t-test. All data presented as mean ± s.e.m.

To identify aggregate-specific growth across the microcolumns, cortical neuronal aggregates were labeled with either green fluorescent protein (GFP) or the red fluorescent protein mCherry via adeno-associated virus 1 (AAV1) transduction (Penn Vector Core, Philadelphia, PA). Briefly, after centrifuging aggregates in the pyramid wells, 1 μL of AAV1 packaged with the human Synapsin-1 promoter was added to the aggregate wells (final titer: ~3×10^10^ viral copies/mL). Aggregates were incubated at 37°C, 5% CO_2_ overnight before the media was replaced twice, after which transduced aggregates were plated in microcolumns as described above, each with one GFP^+^ and one mCherry^+^ aggregate (n = 6). Over multiple DIV, images of the μTENNs were taken on a Nikon A1RSI Laser Scanning confocal microscope paired with NIS Elements AR 4.50.00. Sequential slices of 10-20 μm in the z-plane were acquired for each fluorescent channel. All confocal images presented are maximum intensity projections of the confocal z-slices.

### Viability Assessment

To assess neuronal viability, 5-mm long unidirectional (LE_UNI_) and bidirectional (LE_BI_) constructs and age-matched planar cultures plated on polystyrene were stained with a calcein-AM/ethidium homodimer-1 (EthD-1) assay (ThermoFisher) at 10 and 28 DIV. Metabolically active cells convert the membrane-permeable calcein AM to calcein, which fluoresces green (λexc ~495 nm; λem ~515 nm), while EthD-1 enters membrane-compromised cells and fluoresces red upon binding to nucleic acids (λexc ~495 nm; λem ~635 nm). Briefly, cultures were gently rinsed in DPBS. A solution of calcein-AM (1:2000 dilution; final concentration ~2 μM) and ethidium homodimer-1 (1:500; ~4 μM) in DPBS was added to each culture, followed by incubation at 37°C, 5% CO_2_ for 30 min. Following incubation, cultures were rinsed twice in fresh DPBS and imaged at 10x magnification on a Nikon A1RSI Laser Scanning confocal microscope paired with NIS Elements AR 4.50.00. Viability was quantified as the ratio of the total area of calcein-AM-positive cells to the total area of both calcein-AM-positive and ethidium homodimer-positive cells using ImageJ (National Institutes of Health, MD). Sample sizes for each group were as follows: LE_UNI,5mm_ (n = 4, 4); LE_BI,5mm_ (n = 7, 4); planar cultures (n = 9, 5) for 10 and 28 DIV, respectively. All data presented as mean ± s.e.m.

### Optical Stimulation, Calcium Imaging, and Optical Recording Analysis

To establish whether μTENNs could be coupled with an all-optical system, cortical neuronal aggregates were transduced with either the genetically encoded fluorescent calcium reporter RCaMP1b for optical output or channelrhodopsin-2 (ChR2) for light-based input, via adeno-associated virus 1 (AAV1) transduction as described above (Penn Vector Core). 5-6 mm-long bidirectional μTENNs were then plated with one “input” (ChR2) aggregate and one “output” (RCaMP1b) aggregate at either end (n = 5). ChR2 and RCaMP have been investigated and used for all-optical electrophysiology *in vitro* with minimal spectral overlap, reducing the likelihood of false positive responses due to stimulation of the input aggregate exciting the output aggregate^15^. At 10 DIV, μTENNs were stimulated via an LED optical fiber positioned approximately 1-3mm above the input aggregate, such that the entire aggregate was illuminated. A Plexon Optogenetic Stimulation System with LED modules for each desired wavelength was used to stimulate the μTENNs (Plexon Inc). Stimulation consisted of a train of ten 100ms pulses (1 Hz) at 465nm, within the excitation spectra of ChR2. Each train was repeated three times for a given LED current amplitude (50, 100, 200, 250, 300 mA); amplitudes corresponded to approximate stimulation intensities of 211, 423, 528, and 634 mW/mm^2^ from the tip of the optical fiber and 4.7, 9.3, 18.7, 23.3, and 28.0 mW/mm^2^ at the aggregate, respectively. As a control, μTENNs were stimulated as above at 620nm (outside of the excitation spectra of ChR2) at 300 mA/28.0 mW/mm^2^. Recordings of the μTENNs’ output aggregates were acquired at 25-30 frames per second on a Nikon Eclipse Ti microscope paired with an ANDOR Neo/Zyla camera and Nikon Elements AR 4.50.00 (Nikon Instruments).

To verify whether the fluctuations in calcium reporter fluorescence could be associated with synaptic transmission, bidirectional μTENNs 1.0-1.2mm in length were fabricated and transduced with GCaMP6f (n = 3). At 10 DIV, μTENNs were moved to a stage-mounted warming chamber maintaining incubator conditions (37°C, 5% CO^2^) to record fluorescent calcium reporter activity as described above, with acquisition frequencies of 25-30 frames per second. After 30s of recording spontaneous activity, the NMDA receptor antagonist D-APV (50 μM) and AMPA receptor antagonist CNQX (10 μM) were added to the media containing the μTENNs; recordings were then continued for 20min. Subsequently, media containing D-APV and AMPA was removed and replaced with fresh neuronal culture media. μTENNs were kept at 37°C, 5% CO_2_ overnight, after which spontaneous activity was recorded for an additional 60s.

Following optical stimulation and/or recording, regions of interest (ROIs) containing neurons and background ROIs were identified from the calcium recordings. The mean pixel intensities for each ROI were imported into MATLAB for further analysis via custom scripts (MathWorks Inc). Within MATLAB, the background ROI intensity for each recording was subtracted from active ROIs. Ten such ROIs were randomly selected and averaged to obtain a representative fluorescence intensity trace across each output aggregate. Subsequently, the percent change in fluorescence intensity over time (ΔF/F_o_) was calculated for each mean ROI, where ΔF equals (F_T_ − F_o_), F_T_ is the mean ROI fluorescent intensity at time T, and F_o_ is the average of the lower half of the preceding intensity values within a predetermined sampling window^16^. The peak ΔF/F_o_ for each train was averaged per μTENN for each of the given stimulation intensities. The average maximum ΔF/F_o_ was then compared across stimulation intensities with a one-way ANOVA, with post-hoc analysis performed where necessary with the Tukey procedure (p<0.05 required for significance). Additionally, the peak ΔF/F_o_ of the output aggregate under 620nm/control stimulation was compared to that under 465nm stimulation at 300 mA/28 mW/mm^2^ using an unpaired t-test (p<0.05 required for significance). All data presented as mean ± s.e.m.

### Immunocytochemistry

μTENNs were fixed in 4% formaldehyde for 35 min at 4, 10, and 28 DIV (n = 6, 4, and 8, respectively). μTENNs were then rinsed in 1x PBS and permeabilized with 0.3% Triton X100 + 4% horse serum in PBS for 60 min before being incubated with primary antibodies overnight at 4°C. Primary antibodies were Tuj-1/beta-III tubulin (T8578, 1:500, Sigma-Aldrich) to label axons and synapsin-1 (A6442, 1:500, Invitrogen) to label pre-synaptic specializations. Following primary antibody incubation, μTENNs were rinsed in PBS and incubated with fluorescently labeled secondary antibodies (1:500; sourced from Life Technologies & Invitrogen) for 2h at 18°-24°C. Finally, Hoechst (33342, 1:10,000, ThermoFisher) was added for 10 min at 18°-24°C before rinsing in PBS. μTENNs were imaged on a Nikon A1RSI Laser Scanning confocal microscope paired with NIS Elements AR 4.50.00. Sequential slices of 10-20 μm in the z-plane were acquired for each fluorescent channel. All confocal images presented are maximum intensity projections of the confocal z-slices.

### Cortical Implantation and Intravital Calcium Imaging

As a proof-of-concept for μTENN behavior *in vivo*, bidirectional, approximately 1.5mm-long μTENNs expressing GCaMP were delivered into the brain via stereotaxic microinjection using similar methodology to that described in prior work ^10,11^. Male Sprague-Dawley rats weighing 325-350 grams were anesthetized with isoflurane at 1.0-2.0 liters per minute (induction: 5.0%, maintenance: 1-5-2.0%) and mounted in a stereotactic frame. Meloxicam (2.0 mg/kg) and bupivacaine (2.0 mg/kg) were given subcutaneously at the base of the neck and along the incision line, respectively. The area was shaved and cleaned with betadine solution, after which a small craniotomy was made over the primary visual cortex (V1) (coordinates: −5.0mm AP, ±4.0mm ML relative to bregma). μTENNs were loaded into a needle coupled to a Hamilton syringe and mounted onto a stereotactic arm. To deliver the constructs into the brain without forcible expulsion, the needle was mounted on a micromanipulator and slowly inserted into the cortex to a depth of 1.0 mm such that the dorsal μTENN terminal was left ~500 μm above the brain surface. The plunger of the Hamilton syringe was then immobilized, while the needle containing the μTENN was slowly raised. Upon needle removal from the brain, the dorsal aggregate of the μTENN was immersed in artificial cerebrospinal fluid (aCSF) warmed to 37° C. To protect the dorsal μTENN terminal and enable imaging of the μTENN and surrounding tissue, 2 PDMS discs (5.0mm outer diameter, 2.0mm inner diameter, 0.35mm thickness) were placed over the craniotomy/μTENN and secured to the skull with cyanoacrylate glue. A 3.0mm-diameter glass coverslip was sandwiched between the 2 discs.

At five and ten days post-implant, animals were again anesthetized and mounted on a stereotactic frame for multiphoton calcium imaging of the implant. μTENNs were imaged on a Nikon A1RMP+ multiphoton confocal microscope paired with NIS Elements AR 4.60.00 and a 16x immersion objective. Recordings of the μTENNs’ dorsal aggregates were taken at 3-5 frames per second, similarly to other intravital work^17^. Post-recording, ROIs of μTENN neurons were manually identified, with the mean pixel intensity of each ROI plotted over time. To distinguish neuronal activity from the animal breathing artifact, the fast Fourier transform (FFT) of the mean pixel intensity averaged across 10-15 ROIs was used to identify the frequency peak(s) associated with the observed breathing rate during imaging. Peaks were identified as frequencies whose amplitudes were 2 standard deviations or more than the average amplitude of the Fourier spectra.

### Tissue Harvest and Histology

Post-implant, rats were anesthetized with 150 mg/kg euthasol (Midwest) and transcardially perfused with cold heparinized saline and 10% formalin. After post-fixation of the head overnight, the brain was harvested and stored in PBS to assess μTENN survival and host/μTENN synaptic integration (n = 6). Histology was performed via traditional immunohistology (IHC) and the Visikol clearing method to resolve thicker tissue sections where appropriate.

For traditional IHC, brains were sagittally blocked and cut in 40 μm slices for cryosectioning. For frozen sections, slices were air-dried for 30 minutes, twice treated with ethanol for three minutes, and rehydrated in PBS twice for three minutes. Sections were blocked with 5% normal horse serum (ABC Universal Kit, Vector Labs, cat #PK-6200) in 0.1% Triton-x/PBS for 30-45 minutes. Primary antibodies were applied to the sections in 2% normal horse serum/Optimax buffer for two hours at room temperature. Primary antibodies were chicken anti-MAP2 (1:1000), mouse anti-Tuj1 (1:1000), and mouse anti-synapsin (1:1000). Sections were rinsed with PBS three times for five minutes, after which secondary antibodies (1:1000) were applied in 2% normal horse serum/PBS for one hour at room temperature. Sections were counterstained with DNA-specific fluorescent Hoechst 33342 for ten minutes and then rinsed with PBS. After immunostaining, slides were mounted on glass coverslips with Fluoromount-G mounting media.

In the Visikol method, brains were glued to a vibratome mounting block directly in front of a 5% low EEO agarose post (Sigma A-6103) and placed in PBS surrounded by ice. The brain was cut in 100-200 μm coronal segments with a Leica VT-1000S vibratome until the μTENN implantation site was approximately 1 mm from the cutting face. Subsequently a single 2 mm section containing the microTENN was cut and placed in PBS (frequency setting: 9, speed: 10). The 2 mm brain section was treated at 4°C with 50%, 70%, and 100% tert-butanol, each for 20 minutes. After the ascending tert-butanol steps, the tissue was removed and placed on a kimwipe to carefully remove any excess reagent. Visikol Histo-1 was applied to the sample for 2 hours at 4°C followed by Visikol Histo-2 for 2 hours at 4°C to complete the clearing process. The sample was placed in a petri dish and a hydrophobic well was drawn around the tissue. Fresh Visikol Histo-2 was applied to completely submerge the tissue, which was then covered by a glass coverslip.

Coverslips containing brain slices were imaged on a Nikon A1RMP+ multiphoton confocal microscope paired with NIS Elements AR 4.60.00 and a 16x immersion objective. A 960-nm laser was used to visualize the μTENN containing neurons expressing GFP/GCaMP.

### Data Availability

The data that support the findings of this study are available from the corresponding author upon reasonable request.

## Results

The objectives of our current efforts were threefold: (1) to reproducibly fabricate “living electrode” μTENNs and characterize their growth, architecture, viability, phenotype, and synaptic functionality, (2) to demonstrate the ability to control and monitor μTENNs via light, and (3) to determine whether transplanted μTENN neurons survive *in vivo* and remain viable and active in the host cortex.

### Fabrication and Axonal Outgrowth

In earlier work, μTENNs were seeded with single cell suspensions of primary cortical neurons, which in many cases formed clusters at random sites throughout the microcolumn interior (Figure 1). Current-generation μTENNs were built using cortical aggregates that have been formed prior to plating in the microcolumns, allowing for greater control and reproducibility of the desired cytoarchitecture of discrete somatic and axonal zones (Figure 1). This reproducibility lends itself to consistently creating the desired cytoarchitecture *in vitro*, a necessary step in applying them as living electrodes. Aggregate μTENNs were plated with approximately 8,000-10,000 neurons per aggregate, with microcolumn lengths ranging from 2mm to 9mm; further, both unidirectional (with one aggregate) and bidirectional (with two aggregates at either end) μTENNs were plated with different lengths. Healthy axonal outgrowth was found across all aggregate μTENNs along the ECM core within the first few days *in vitro* through analysis of phase microscopy images (Figure 2). All aggregate μTENNs exhibited rapid axonal growth rates, peaking at 1101.8±81.1 microns/day within the LE_BI,9_ group. In general, aggregate μTENNs displayed maximal growth at 3 days *in vitro* (DIV); exceptions included LE_BI,8_ and LE_UNI,5_, where the fastest growth was observed at 5 DIV, and LE_UNI,2_, with an average maximum growth rate of 580±43.9 microns/day at 1 DIV (Figure 2F, Table 1). Comparatively, dissociated μTENNs exhibited a peak growth rate of 61.7±5 microns/day at 1 DIV, representing a nearly 17-fold reduction relative to the aggregate μTENNs within the LE_BI,9_ group (Figure 2F, Table 1). Planar control cultures exhibited an average growth rate of 38.1±19.4 microns/day from 1 to 3 DIV and 39.1±20.6 microns/day from 3 to 5 DIV, after which single neurites could not be identified (Table 1). The two planar growth rates did not differ significantly, yielding a cumulative average growth rate of 38.6±20.0 microns/day.

One-way ANOVA of the average maximum growth rate identified a significant main effect of the LE group (F-statistic = 14.1, p < 0.0001). Subsequent Bonferroni analysis on pairwise comparisons revealed that the average maximum growth rates of bidirectional μTENNs ranging from 7 to 9 mm in length (LE_BI,7_, LE_BI,9_) were statistically higher than those 2 to 3 mm in length (LE_BI,2_, LE_BI,3_) (p < 0.001); additionally, the maximum growth rate of LE_BI,9_ was found statistically higher than that of LE_UNI,2_ (p < 0.01) and LE_UNI,5_ (p < 0.05) (Figure 2). ANOVA of the minimum growth rate did not detect any differences across LE groups (F-statistic = 1.17, p = 0.332), while ANOVA of the average crossing time (F-statistic = 12.99, p <0.0001) and Bonferroni post-hoc analysis showed that LE_BI,7_ and LE_BI,9_ axons crossed the length of the microcolumn later than those of LE_UNI,2_, LE_BI,2_, LE_BI,3_, and LE_BI,5_ (Figure 2E). μTENNs within LE_UNI,5_ did not, on average, fully span the construct length by 10 DIV (Figure 2, Table 1).

**Figure 2:**
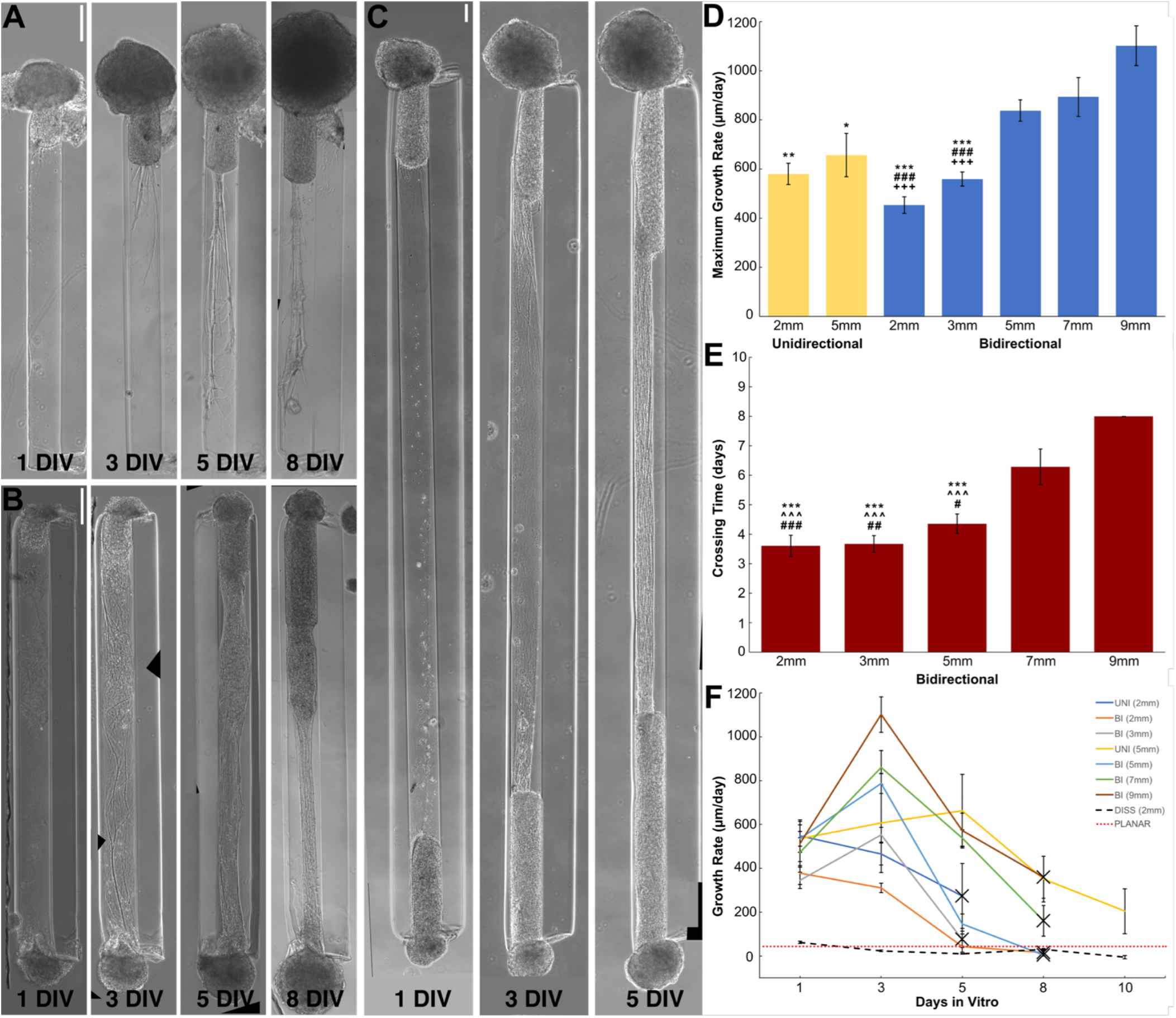
Axonal Growth in Aggregate μTENNs Over Time. Both unidirectional **(A)** and bidirectional **(B)** μTENNs displayed robust axonal outgrowth along the ECM core. While outgrowth within unidirectional μTENNs peaked within the first 3 DIV before declining, bidirectional μTENN axons crossed the length of the microcolumn, synapsing with the opposing aggregate by 5 DIV. Representative 2mm μTENNs shown at 1, 3, 5, and 8 DIV. **(C)** Longer bidirectional μTENNs (5 mm) took more time to develop, but still showed robust growth. Representative 5mm μTENN shown at 1, 3, and 5 DIV. **(D)** Average maximum growth rates across μTENN groups. Symbols denote significant differences vs. 9mm bidirectional (*), 7mm bidirectional (#), and 5mm bidirectional (+) μTENNs, respectively. Symbol count denotes significance level (1: p < 0.05; 2: p < 0.01; 3: p < 0.001). **(E)** Average crossing times across μTENN groups. 5mm unidirectional μTENNs did not fully cross by 10 DIV and were not included. Symbols and symbol counts match those described in (D), with the addition of significance vs. 8mm bidirectional (^). **(F)** Growth rates for unidirectional, bidirectional, and dissociated/traditional μTENNs at 1, 3, 5, 8, and 10 DIV; dashed red line represents the average growth rate for planar cultures. Growth rates were quantified by identifying the longest neurite from an aggregate in phase microscopy images (10X magnification) at the listed timepoints. Crosses indicate axons crossing the length of the microcolumn (unidirectional) or connecting between aggregates (bidirectional). Error bars denote s.e.m. Scale bars: 200 μm.

**Table 1:** μTENN Growth Characterization. Data presented as mean ± s.e.m. in units of microns/day (Initial, Maximum, and Minimum Growth Rates) and days in vitro (Crossing Time). LE subscripts indicate unidirectional (UNI), bidirectional (BI), or dissociated (DISS) μTENNs and the microcolumn length in millimeters.

| Growth Rates | $LE_{UNI,2}$ | $LE_{BI,2}$ | $LE_{BI,3}$ | $LE_{UNI,5}$ | $LE_{BI,5}$ | $LE_{BI,7}$ | $LE_{BI,9}$ | $LE_{DISS,2}$ | Planar |
| --- | --- | --- | --- | --- | --- | --- | --- | --- | --- |
| Initial | $547 \pm 73.2$ | $378 \pm 51.9$ | $345 \pm 35.4$ | $525 \pm 21.9$ | $535 \pm 62.9$ | $472 \pm 66.9$ | $513 \pm 100.7$ | $61.7 \pm 5.01$ | $38.1 \pm 19.4$ |
| Maximum | $580 \pm 43.9$ | $453 \pm 33.8$ | $559 \pm 28.7$ | $656 \pm 88.3$ | $838 \pm 43.2$ | $894 \pm 79.2$ | $1102 \pm 81.1$ | $61.7 \pm 5.01$ | $39.1 \pm 20.6$ |
| Minimum | $430 \pm 73.0$ | $248 \pm 22.7$ | $324 \pm 38.0$ | $202 \pm 74.9$ | $395 \pm 56.4$ | $336 \pm 57.4$ | $313 \pm 68.3$ | $-5.37 \pm 7.50$ | $38.1 \pm 19.4$ |
| Crossing Time | 4.33 | 3.60 | 3.67 | NA | 4.35 | 6.29 | 8.0 | N/A | N/A |

### μTENN Viability

Neuronal survival was quantified via live/dead staining and confocal microscopy for short unidirectional and short bidirectional μTENNs at 10 and 28 DIV (Figure 3). Age-matched planar cultures served as controls. Percent viability was defined as the ratio of the summed area of calcein-AM-positive cells to that of all stained cells (i.e. both calcein-AM^+^ and ethidium homodimer^+^ cells). Neuronal survival in μTENNs was observed to persist up to at least 28 DIV, with further demonstration of survival out to 40 DIV (Figure 3). ANOVA showed that although the DIV was a significant main effect (F-statistic = 32.21, p < 0.0001), the LE/culture group was not a significant factor (p > 0.84). The interaction effect was significant (p < 0.01), so Bonferroni analysis was used to compare groups at each time point (Figure 3G). Viability of planar cultures at 28 DIV (53.6%) was found statistically lower than that of LE_UNI_ (80.3%) (p < 0.05), LE_BI_ (84.8%) (p < 0.001), and planar cultures (97.7%) (p < 0.0001) at 10 DIV. Moreover, planar culture viability at 10 DIV surpassed those of both LE_UNI_ (68.1%) and LE_BI_ (69.0%) at 28 DIV (p < 0.01). Overall, planar cultures exhibited a 45% decline in viability from 10 to 28 DIV, while LE_UNI_ and LE_BI_ showed a 15.2% and 18.6% drop over time, respectively (Figure 3).

**Figure 3:**
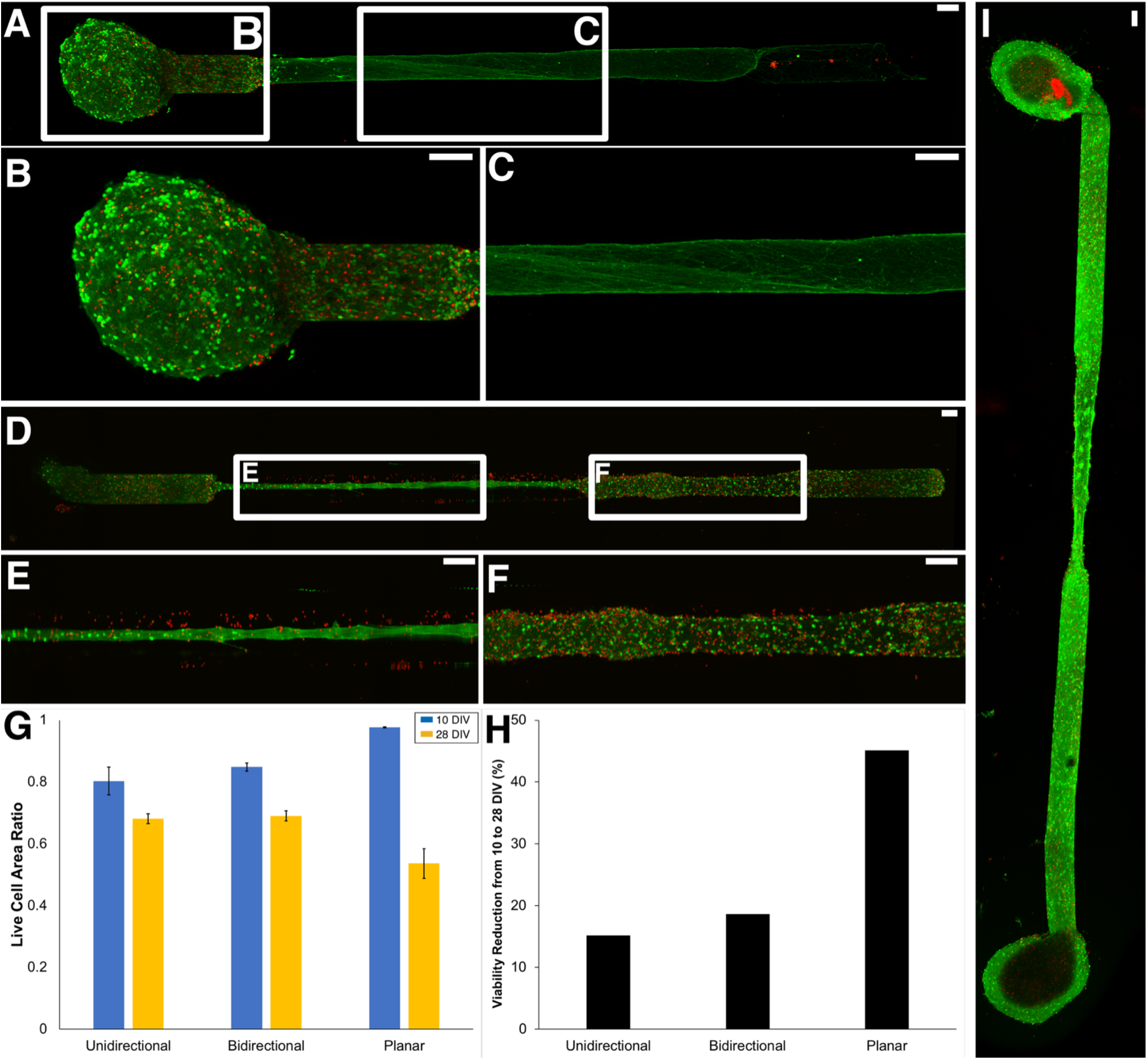
μTENN Viability. Viability for unidirectional and bidirectional μTENNs and age-matched two-dimensional controls was quantified via live-dead (calcein-AM/ethidium homodimer) staining at 10 and 28 DIV. **(a, b, c)** Representative confocal live-dead images showing live cells (green), dead cells (red), and an overlay of a unidirectional μTENN at 10 DIV, with outlined insets below. **(d, e, f)** Representative confocal live-dead image of a bidirectional μTENN at 28 DIV, with outlined insets below. **(G)** The average proportion of live to total (live + dead) cell body area for each experimental group and timepoint. Two-way ANOVA and post-hoc analysis revealed several statistically relevant pairwise differences (* = p < 0.05; ** = p < 0.01; *** = p < .001). Symbols denote significant differences vs. planar cultures at 10 DIV (#) and 28 DIV (*). Error bars denote s.e.m. Sample sizes: n = 4 and 4 (unidirectional); 7 and 4 (bidirectional); 9 and 5 (controls) for 10 and 28 DIV, respectively. **(H)** The percent change in viability across experimental groups. All groups showed a decline in viability, with the planar cultures nearing a three-fold drop in viability relative to the μTENNs. **(I)** Live-dead stain of a μTENN at 40 DIV. Scale bars: 100 μm.

### μTENN Architecture and Synaptogenesis

To characterize μTENN architecture, bidirectional μTENNs were either labeled with GFP and mCherry and imaged over time or fixed and immunolabeled at set timepoints to identify cell nuclei, axons, and synapses (Figure 4). Confocal images of GFP/mCherry μTENNs revealed that upon making contact with opposing axons, projections continued to grow along each other towards the opposing aggregate, confirming physical interaction and integration between the two neuronal populations (Figure 4). Immunolabeling revealed that neuronal somata were localized almost exclusively to the aggregates, which were spanned by long axons, as indicated with Tuj-1 (Figure 4H); axons and dendrites were also found within the aggregates from intra-aggregate connections, presumably formed upon or shortly after plating. Synapse presence was qualitatively assessed using the sum area of synapsin^+^ puncta across the specified timepoints. A modest distribution of synapsin within μTENN aggregates was observed, as well as an increase in synapsin expression within the lumen of the microcolumns, suggesting that neurons within bidirectional μTENNs synaptically integrate and therefore have the capacity to communicate between aggregates.

**Figure 4:**
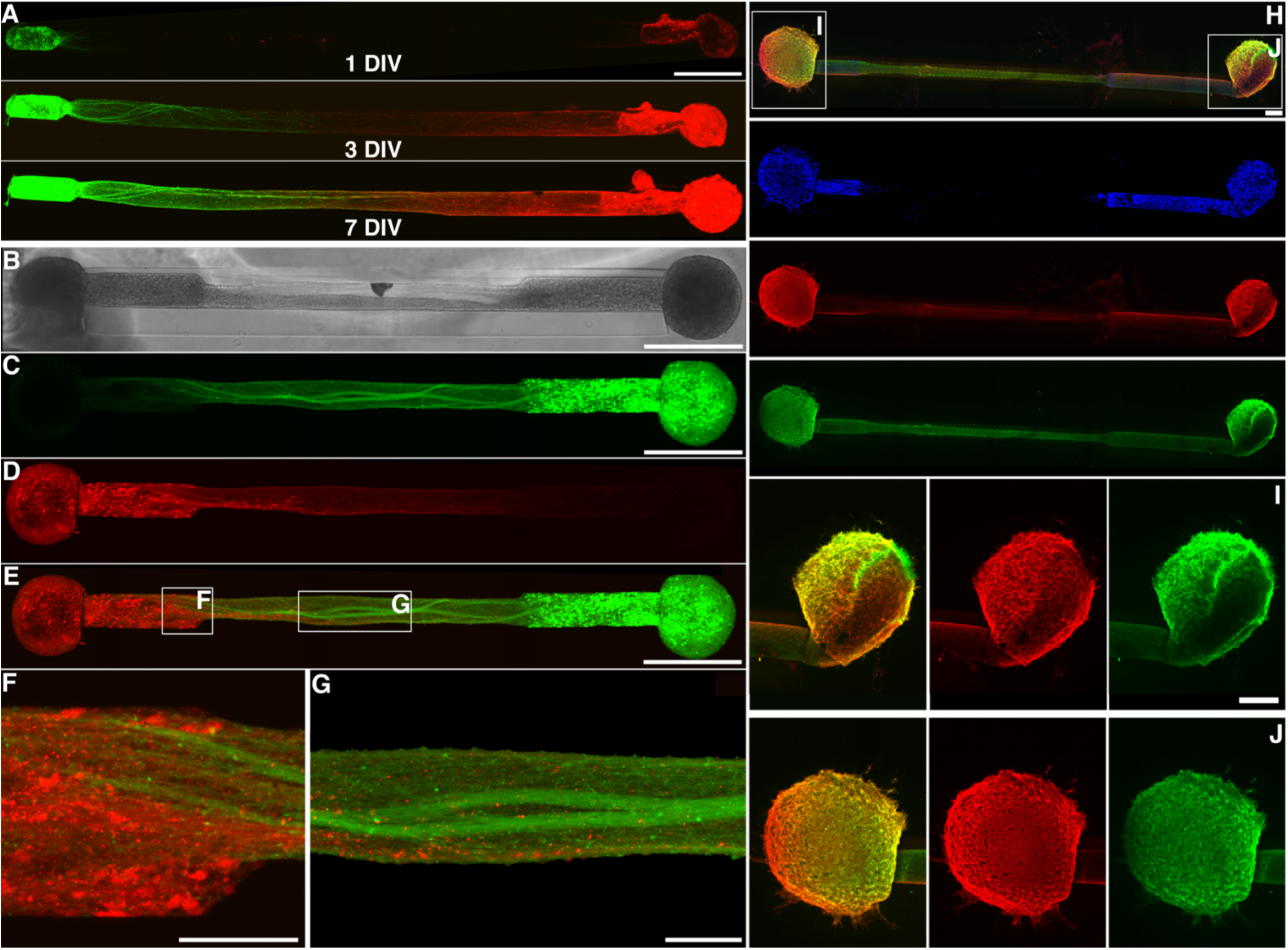
μTENN Growth and Architecture. Bidirectional μTENNs were labeled with GFP (green) and mCherry (red) to observe aggregate-specific axonal growth and structure *in vitro*. **(A)** Confocal reconstructions of a bidirectional, GFP/mCherry-labeled μTENN at 1, 3 and 7 DIV. **(B)** Phase image of a bidirectional, GFP/mCherry-labeled μTENN at 5 DIV. **(C-E)** Confocal reconstruction of the μTENN from (B) at 7 DIV, with insets showing axons from each aggregate growing along each other **(F)** and axons from one aggregate making contact with the opposite population **(G)**. **(H)** Confocal reconstruction of a representative bidirectional μTENNs at 10 DIV immunolabeled for cell nuclei (Hoechst; blue), axons (Tuj-1; red), and synapses (synapsin; green). Insets in (H) refer to callout boxes **(I)** and **(J)** showing zoom-ins of synapses, axonal networks, and the overlay of the two. Scale bars: 500 μm (A, B, C, E); 100 μm (F, G); 200 μm (H, I).

### Calcium Imaging and Optical Stimulation

Bidirectional μTENNs expressing the calcium reporter GCaMP6f exhibited spontaneous oscillations in the delta band (1-5 Hz) in the absence of external stimulation, with the synchronicity of oscillation between aggregates suggesting the potential formation of synaptic networks. Moreover, the introduction of the NMDA and AMPA receptor antagonists D-APV and CNQX to media containing bidirectional μTENNs reversibly abolished endogenous activity as measured by the calcium reporter GCaMP6f (data not shown), indicating that the calcium transients observed may reflect action potential firing due to synaptic transmission. Bidirectional μTENNs were also engineered to enable light-based stimulation and concurrent calcium imaging *in vitro* by transducing one aggregate with ChR2 and the opposing aggregate with RCaMP. Upon illumination of ChR2^+^ (input) aggregates with 465nm light (stimulation wavelength of ChR2), the opposing RCaMP^+^ (output) aggregates exhibited timed changes in fluorescence intensity in response. As a negative control, the input aggregate was exposed to 620nm light (off-target wavelength), revealing no readily observable responses; the mean peak ΔF/F_o_ of the output aggregate was significantly greater under 465nm stimulation than 620nm stimulation at 634mW/mm^2^ (p < 0.05). Collectively, these findings suggest that the changes in ΔF/F_o_ under 465nm stimulation reflected synaptically mediated firing of neurons in the output aggregate in response to light-based activation of neurons within the input aggregate. Although there was high variability in ΔF/F_o_ between μTENNs, the percent change relative to baseline fluorescence due to optical stimulation could be reproducibly distinguished from endogenous activity across all the μTENNs studied and the average maximum ΔF/F_o_ positively correlated with the stimulation intensity (Figure 5). Overall, these results suggest that light-based stimulation of the input aggregate resulted in controllable signal propagation and modulation of activity in the output aggregate.

**Figure 5:**
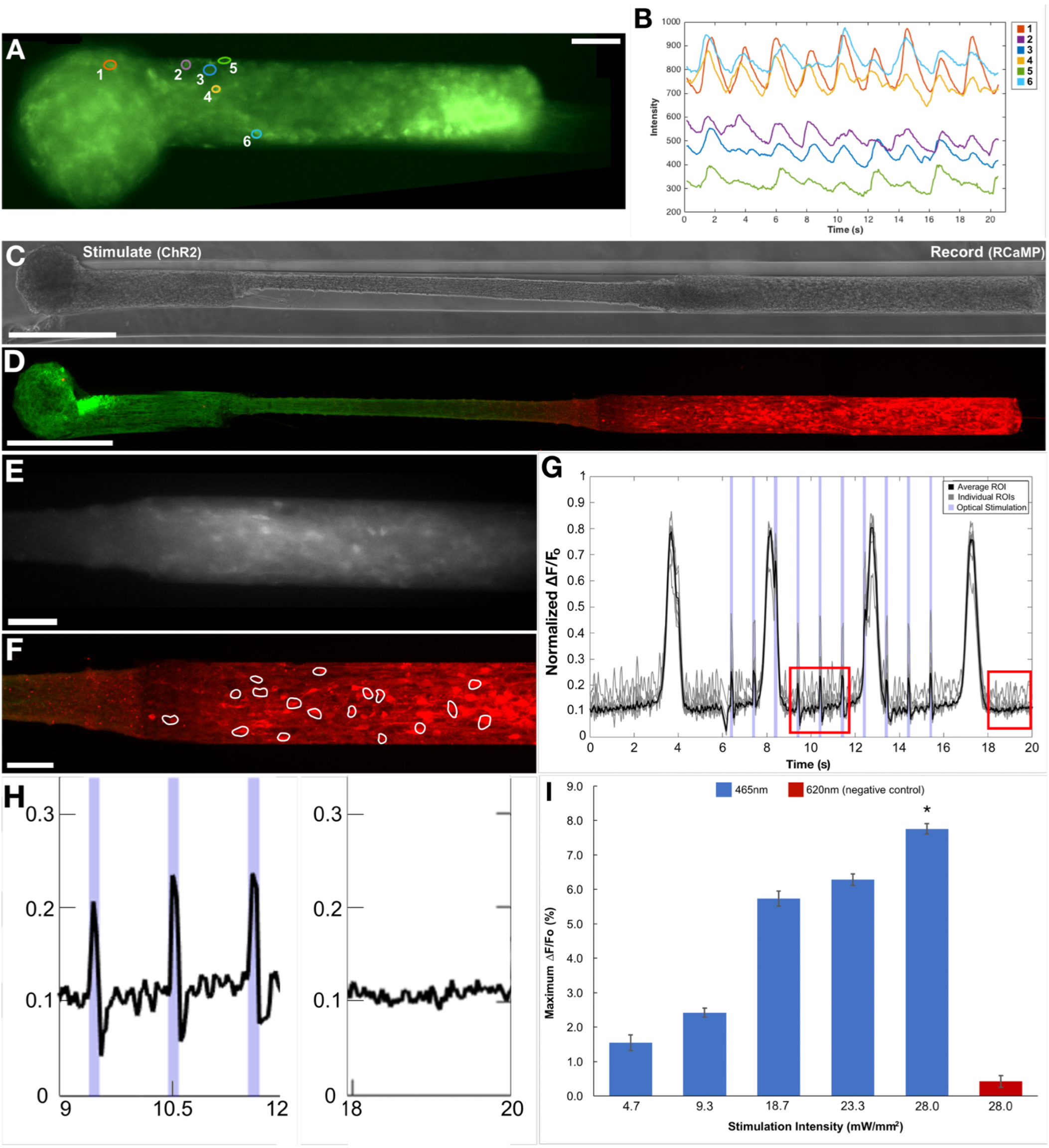
Simultaneous Optical Stimulation and Recording in μTENNs. μTENNs transduced to express both an optical actuator and a fluorescent calcium reporter may be controlled and monitored with light. **(A)** Representative micro-TENN transduced with GCaMP6 at 10 DIV. **(B) I**ntensities of the ROIs from (A) recorded over time. Intensities shown are normalized to the average of a background region (i.e. away from the micro-TENN). **(C)** Phase image and **(D)** confocal reconstruction of a μTENN at 10 DIV *in vitro*, with the left aggregate transduced with ChR2 and the right aggregate transduced with the calcium reporter RCaMP. **(E)** The RCaMP-positive aggregate from (D) under fluorescent microscopy during recording (16 fps). **(F)** Confocal reconstruction of (D) post-stimulation. ROIs containing single neurons were manually defined (white outlines). **(G)** Normalized pixel intensity of ROIs within the RCaMP+ aggregate from (a-c) during stimulation. Grey lines indicate representative, user-defined ROIs randomly selected for analysis, which were averaged to obtain a mean ROI of the aggregate (solid black line). The timestamps of a single train of 1 Hz, 100ms stimulation pulses are shown as blue bands along the abscissa. The changes in pixel intensity due to stimulation of the input aggregate can be seen as sharp spikes occurring within the endogenous, large-amplitude slow-wave activity. **(H)** Zoom-ins of the red insets from (G) showing μTENN activity during (left) stimulation and after (right) optical stimulation. **(I)** Average maximum ΔF/F_o_ across stimulation intensities (at the aggregate). Although the maximum ΔF/F_o_ trended upward, the differences were not significant across intensities. Statistical comparison revealed that stimulation with the control wavelength (620nm) yielded significantly lower maximum ΔF/F_o_ than with 465nm (* = p < 0.05). Scale bars: 100 μm.

### Implantation and Intravital Calcium Imaging

μTENNs – fabricated as described above and transduced to express GCaMP6 – were implanted as a proof-of-concept for living electrode survival, integration, and function. One week and one month-post injection in the rodent brain, constructs were found to have survived and maintained the preformed somatic-axonal architecture, with cell bodies predominantly localized to one or both microcolumn terminals and spanned by axonal tracts (Figure 6). Large, dense clusters of GCaMP^+^ cell bodies (aggregates) were found at the dorsal and ventral regions of implantation, with axons and dendrites within the lumen spanning the two locations (Figure 6). There was also significant neurite outgrowth from the ventral end of the living electrode, with structural evidence of synapse formation with host neurons (Figure 6). In some cases, there was also neuronal migration up to several millimeters from the ventral implant location, although the presence and extent of migration varied across implants. Multiphoton imaging revealed GCaMP-positive μTENN neurons in V1 at both 5 and 10 days post-implant (Figure 7). The breathing of the anesthetized animal was controlled via monitoring and controlled isoflurane delivery, and changes in GCaMP fluorescence intensity due to breathing artifact were readily identified within the FFT of the time-lapse recordings as a ~0.5-0.7 Hz peak (Figure 7). Non-artifact changes in GCaMP intensity were present in the delta band (1-5 Hz), indicating μTENN survival and neuronal activity (Figure 7). Putative activity was also present at frequencies below 1 Hz, within the reported range reported for slow-wave cortical activity under anesthesia and during sleep^18,19^. Calcium recordings of μTENNs at both 5 and 10 days post-implant reflected those of non-implanted μTENNs at 10 DIV, which was also dominated by low frequency activity in the 1-5 Hz range (as shown in Figure 5).

**Figure 6:**
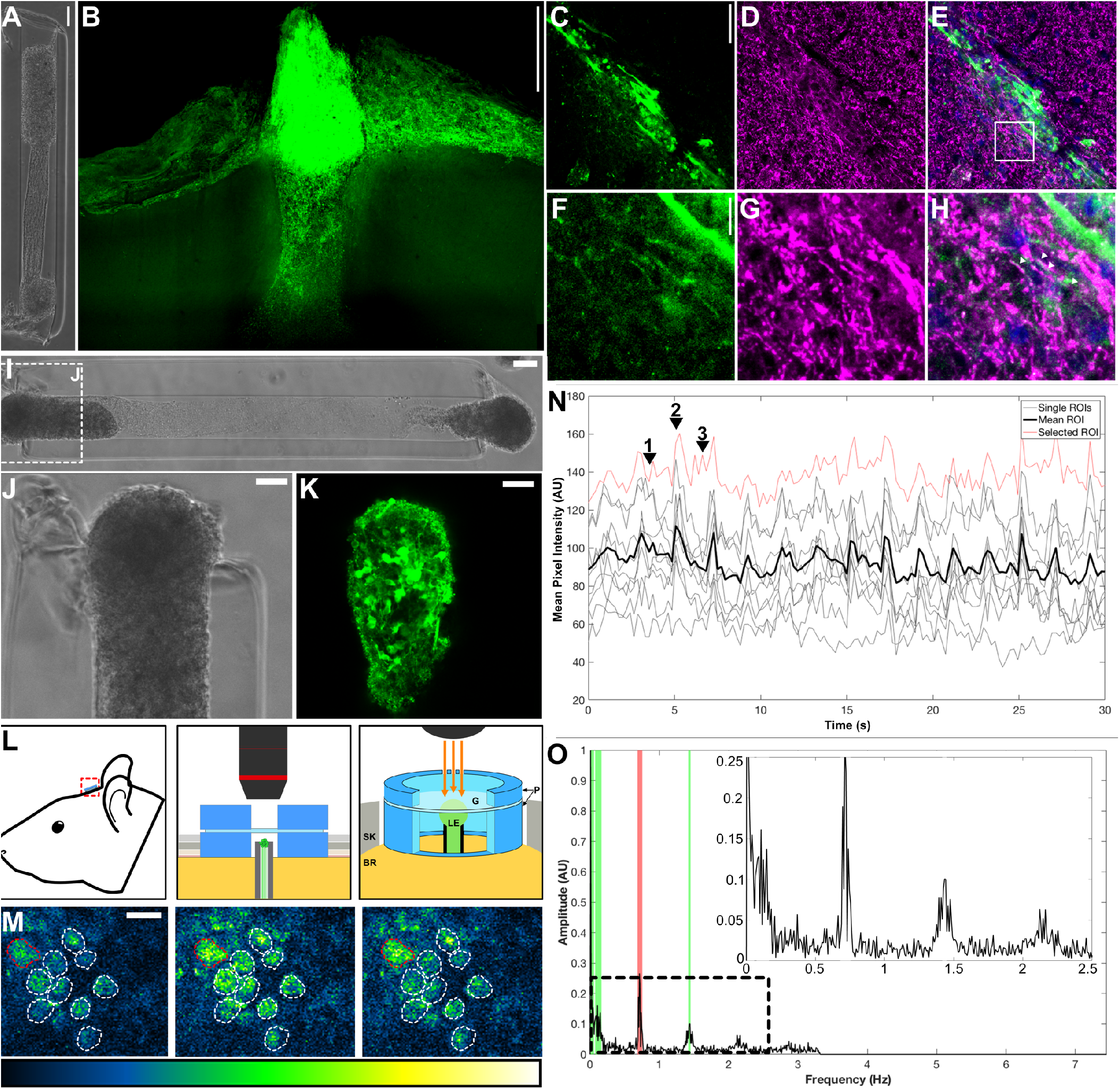
Living Electrode Survival, Integration, and Function *in Vivo*. **(A)** Phase image of a bidirectional μTENN prior to implantation; aggregates have been internalized to the microcolumn. **(B)** Multiphoton image of the μTENN from (A) at one-month post-implant, showing GCaMP-positive μTENN neurons and processes within and immediately surrounding the construct. At one month, the dorsal aggregate had descended into the microcolumn, suggesting externalized aggregates may be required to maintain a cohesive neuronal population at the surface. **(C-E)** Confocal image of a μTENN at one-month post implant, with synapsin^+^ puncta at and around the μTENN/brain interface. Shown are μTENN neurons (GFP; green), synapses (synapsin; purple) and nuclei (Hoechst, blue). **(F-H)** Zoom-ins of inset from (E) with colocalization of synapsin and GFP suggesting synaptic integration. **(I)** Phase image of a GCaMP+ μTENN prior to implant. Inset refers to **(J)** showing the dorsal aggregate. **(K)** Multiphoton image of the dorsal aggregate of the μTENN, acquired immediately post-implant. **(L)** Conceptual schematic of the μTENN and cranial window. Inset shows in more detail the PDMS rings (P) sized to the skull craniotomy (SK) securing the glass coverslip (G) and protecting the implanted μTENN and underlying brain (BR), which may then be imaged chronically (orange arrows). **(M)** Single frames from multiphoton recording of the μTENN from (a-b) at 10 days post-implant during low activity (left), breathing (middle), and non-artifact neuronal activity (right). ROIs approximating single neurons are outlined. The LUTs scale (0-4096) is provided below. **(N)** Time course of calcium fluorescence from (E), showing the individual ROIs (grey/red) and average fluorescence across the aggregate. The red trace represents the ROI outlined in red in (E). Numbered arrows denote timestamps from (E). A sample of the recording can be found in Supplemental Video 3. **(O)** Fourier transform of the data from (F), showing spectral peaks due to the breathing rate as measured during imaging (red) and neuronal activity (green). Inset shows low-frequency activity similar to that observed *in vitro*. Scale bars: 500 μm (B); 100 μm (A); 50 μm (C); 10 μm (F); 100 μm (I); 50 μm (J, K); 20 μm (M).

## Discussion

Microelectrodes—the current gold standard for recordings—have been deployed successfully on the order of months, and less frequently years, in rodents, non-human primates, and human patients^1,20–22^. However, microelectrode-based BCIs generally succumb to a complex combination of abiotic and biological factors, including neuronal loss/migration, gliosis, biofouling, electrode movement, and/or mechanical failure – which impede stability, specificity, and clinical deployment^1–5,23^. Optogenetic strategies for neuromodulation permit more selective stimulation, but must address formidable challenges such as restricting the vector of interest to targeted cells, addressing the scattering and limited tissue penetration of light, and activating transduced cells without overheating brain tissue^24–27^. Efforts to minimize inflammation have yielded more compliant electrodes and electrode coatings/co-factors; however, the chronic foreign body response, consequent signal drop, and increase in stimulation thresholds continue to affect many current systems.

As an alternative to conventional microelectrodes and/or optogenetics strategies, μTENNs as living electrodes may present a neuromodulation/recording platform with improved selectivity and longevity. By being fully fabricated *in vitro*, μTENNs leverage advantages of optogenetics while (1) avoiding any inherent risks of introducing active viruses *in vivo*, (2) restricting viral expression to the μTENN neurons only, and (3) leveraging well-established stereotactic neurosurgical techniques. The microcolumn size may be minimized to reduce the microinjection footprint, while its material properties (e.g. stiffness) and potential co-factors (e.g. anti-inflammatory/growth factor release) may be tailored against any subsequent foreign body response. Moreover, while the constructs in this study were predominantly glutamatergic, μTENNs may be seeded with other neuronal subtypes for various applications (e.g. inhibitory or dopaminergic neurons) to enable more targeted integration based on the synaptogenetic behaviors of the subtype. Finally, as the μTENN-brain interface is synaptic, living electrodes may potentially remain stable in the brain for extended periods of time. Given the potential benefits of the μTENN paradigm, the work described here represents a critical foundation in developing these implantable, engineered neural networks into a viable neural interface. Indeed, the biofabrication of phenotypically-controlled, fully-implantable axon-based living electrodes may provide a useful tool for the neuroscience community to probe and modulate deep neural circuitry based on biological specificity provided by natural, synaptic inputs while being accessible on the brain surface for optical read-out/control.

A key objective of the current work was the development of advanced methodology for the biofabrication of fully implantable, three-dimensional (3D) cylindrical microtissue replicating key neuroanatomical features: discrete populations(s) of phenotypically controlled neurons spanned by dense bundles of longitudinally aligned axonal tracts. Remarkably, we identified biofabrication techniques that not only consistently created our desired cytoarchitecture, but also resulted in the emergence of accelerated axonal outgrowth and improved neuronal survival versus traditional, planar cultures. In particular, we found that neuronal aggregate-based biofabrication allowed for more standardized construct architecture and repeatable studies compared to single-cell suspensions. Notably, we found that aggregate-based μTENNs exhibited faster axonal growth and greater total axonal lengths than their dissociated counterparts. The observed growth rates for dissociated μTENNs were similar to those in planar cultures, which averaged nearly 40 μm/day over the first 3 days (Figure 2). This falls within the growth reported in literature for cortical axons, which have reached lengths of up to 100-1000 μm over 3 days in planar cultures^28,29^. However, the peak axonal growth rates that were measured from aggregates greatly exceeded those in planar counterparts and dissociated μTENNs by 2 orders of magnitude, or over 1000 μm/day. Although further investigation is needed, we have identified a few potential causes of this significant benchmark. First, the restriction of axonal outgrowth to the microcolumn interior resulted in the formation of “bundles” of axons from the aggregates, which may be directionally self-reinforcing and accelerate linear extension. Second, the lack of synaptic targets within the microcolumn may reduce axon branching between aggregates, which would otherwise slow growth cone movement^28,30–33^. Further, although longer μTENNs generally exhibited faster growth than shorter ones, initial growth rates did not vary significantly across different lengths. Thus greater separation between the aggregates may be necessary to establish either sufficient chemotactic gradients or a “ramp up” of growth machinery, such that maximal growth rates are only reached when targets are several mm away. Finally, axon growth was mediated, if not accelerated, by the collagen-laminin ECM, as its constituents are known to support axonal growth^34^. Indeed, aggregate μTENNs created without ECM had limited neurite outgrowth and did not develop inter-aggregate connections (data not shown).

μTENN neuronal viability was shown to persist for up to 40 DIV, suggesting their potential for use in long-term *in vitro* studies. Interestingly, the decline in viability from 10 to 28 DIV was lower for both unidirectional and bidirectional μTENNs than for planar cultures. While the cause of this improved survival potential has not been fully investigated, established work suggests that neurons exhibit better growth and survival in 3D environments, which more accurately approximate conditions *in vivo^35^*. Similarly, the anatomically inspired 3D microstructure of the neuronal aggregates and axonal bundles may enable neurons to better self-regulate and remain healthy compared to 2D cultures.

In addition to rapid growth and improved survival, a key outcome was the determination of functional connectivity across the neuronal populations mediated by the engineered axonal tracts. Here, structural evidence of neuritic and synaptic integration was visualized as early as 4 DIV within the microcolumns. As the primary points of contact and communication between neurons, synapses are often used to determine the functional maturity of neuronal cultures^36,37^. Synapsin^+^ puncta were observed to increase between 4 DIV and 28 DIV, suggesting that μTENN neurons form functional connections soon after plating which mature and expand over time, consistent with literature for planar cortical cultures^37^. Future network connectivity studies may more fully characterize the development and distribution of intra-versus inter-aggregate synapses; however, it is likely that intra-aggregate synapses initially dominate the total synapse population before axons span the aggregates and enable inter-aggregate synapses to form. These analyses would build on the aggregate-specific labeling achieved here to distinguish structures from either aggregate (Figure 4). Overall, these results indicate that μTENNs are capable of quickly and consistently forming the desired μTENN architecture – important for the biofabrication and scale-up of experimentally useful constructs – which is maintained over weeks *in vitro*. Moreover, the μTENNs’ structure may make them an ideal system for studying neuronal growth, maturation, and network dynamics *in vitro*, with characteristics approximating the 3D architecture of connectome-spanning structures in the mammalian brain more closely than planar cultures^35^.

Initially, we measured spontaneous activity in and across the aggregates (Figure 5, Supplemental Movies 1 & 2), which consisted primarily of delta oscillations (1-5 Hz). Concurrent network analyses have shown that μTENNs *in vitro* exhibit inter-aggregate synchronicity within the delta band^38^. The introduction of glutamatergic receptor blockers and subsequent suppression of GCaMP^+^ activity implicate synaptic transmission as the primary contributor to changes in reporter fluorescence. Optical stimulation and recordings of evoked activity across aggregates further demonstrate the presence of functional axonal tracts and synaptic-mediated integration across two aggregate populations. These important steps validated the functionality and long-distance transmission across the axonal tracts within the microcolumns and, crucially, demonstrate the ability of these constructs to serve as an “all optical” input-output platform for experimental use *in vitro* and/or for circuit modulation *in vivo*. Indeed, post-transplant into the rat cortex, we found that this activity persisted, along with slow-wave activity recorded below 1 Hz. Slow-wave oscillations have been recorded under anesthesia and during slow-wave sleep, as well as in cortical neuronal cultures *in vitro*^18, 19^. Whether the <1 Hz activity observed within the μTENNs reflects cortical activity is unknown at present and will be further determined through continued intravital, functional, and histological analyses at longer timepoints. Combined with the presented histology, there is strong evidence that μTENNs survive post-transplant and form putative synapses with the cortex, although we observed significant overgrowth and integration with a subset of our transplants. As such, controllability over neuronal migration and the targeting of synaptic integration remains an ongoing design challenge that will need to be addressed to ensure proper function. This may be done by controlling neuronal subtype as discussed, or by otherwise manipulating the transplant environment to promote more targeted integration, e.g. introducing or promoting expression of trophic factors implicated in axonal guidance and/or synaptic pruning during development^39,40^. Potential physical targeting methods include a porous membrane at the ventral terminal to restrict neuronal migration while permitting axonal projections between the μTENN and host brain.

In summary, we have created so-called living electrodes – cylindrical hydrogel-encapsulated neuronal populations linked by functional axonal tracts – and demonstrated their biofabrication, functional validation, targeted delivery, and survival and integration post-transplant. These milestones lay the groundwork for more in-depth investigations of the translational utility of the μTENNs following targeted transplant in the cerebral cortex or other anatomical targets. Future work will assess the ability of transplanted μTENNs as an experimental tool to modulate (input) and/or record (output) brain activity as a neural interface (Figure 1). For inputs, unidirectional, optogenetically-active μTENNs may bypass the light scattering and limited penetration depth of conventional optogenetic methods by relaying light stimulation at the cortical surface into synaptic inputs to the desired target. For outputs, bidirectional μTENNs may be transduced with GCaMP^+^ or similar reporters and transplanted. Upon forming synapses with host neurons, GCaMP^+^ μTENNs may be used to monitor neuronal activity deeper in the brain, providing actionable representations of deeper neural signals to the brain surface. Taken together, these results serve as an early proof-of-concept for μTENNs as a platform for biologically-based neuromodulation. Through optogenetic and tissue-engineering techniques, we have advanced the development of preformed, implantable neural networks as a potentially long-term neural interface at the intersection of neuroscience and engineering.

## Author Contributions

Conceptualization: D.K.C., J.A.W., M.D.S., H.I.C.; Methodology: D.K.C., D.O.A., L.A.S., J.P.H., A.D.N., J.C.B., D.P., H.I.C., J.A.W.; Formal Analysis: D.O.A.; Investigation: D.O.A., J.C.B.; Resources: R.H.K.; Visualization: D.O.A.; Writing – Original Draft: D.O.A.; Writing – Review & Editing: D.O.A., D.K.C., L.A.S., J.P.H., A.D.N., J.C.B., D.P., R.H.K., H.I.C., J.A.W., M.D.S; Supervision: D.K.C., J.A.W., M.D.S., R.H.K., H.I.C.; Project Administration: D.K.C.; Funding Acquisition (primary): D.K.C.

**Supplemental Videos 1 & 2: Calcium Imaging within μTENNs.**

The two videos show spontaneous network activity within μTENN aggregates, visualized with the genetically encoded calcium reporter GCaMP6f. Video 1 shows the same μTENN aggregate shown in Figure 6E. Video 2 shows a bidirectional μTENN approximately 1.1mm in length, imaged at 10 days in vitro. Note that GCaMP activity can be seen in both the aggregate and axonal regions of the μTENN.

**Supplemental Video 3: Simultaneous Optical Stimulation and Recording.**

This video shows the RCaMP^+^ aggregate of a ChR2/RCaMP μTENN approximately 6mm in length, imaged at 10 DIV during an optical stimulation/recording experiment. Red arrows indicate optical stimulation of the ChR2^+^ aggregate (outside of the field of view) at 470 nm. Output power at the optical fiber for the pulse trains was 106, 211, 317, 423, 528, and 634 mW/mm^2^; corresponding to 50, 100, 150, 200, 250, and 300 mA current amplitude, respectively. An increase in the RCaMP^+^ aggregate fluorescence can be seen as the intensity of the pulse trains increases over time.

